# Treatment of non-alcoholic steatohepatitis with recombinant Orosomucoid 2, an acute phase protein that attenuates the Erk1/2-PPARγ-Cd36 signaling pathway

**DOI:** 10.1101/2022.10.22.513320

**Authors:** Li Li, Haoming Sun, Jionghao Chen, Cong Ding, Xiaojun Yang, Hua Han, Qingzhu Sun

**Affiliations:** Department of Animal Science, College of Animal Science and Technology, Northwest A&F University, Yangling, Shaanxi, China; Department of Biomedicine, Future Agriculture Institute, Northwest A&F University, Yangling, Shaanxi, China

**Keywords:** Orm2, NAFLD, recombinant protein, fatty acid uptake, Cd36

## Abstract

Therapeutic administration of recombinant proteins has been used for the treatment of various diseases in multiple studies. Herein, we investigated the function of the acute phase protein Orosomucoid 2 (Orm2), which is mainly secreted by hepatocytes, and evaluated its potential as a therapeutic strategy for non-alcoholic fatty liver disease (NAFLD) and non-alcoholic steatohepatitis (NASH). We here revealed that a high expression of Orm2 protected mice from high-fat diet (HFD)-induced obesity. The pharmacological administration of recombinant ORM2 protein attenuated hepatic steatosis, inflammation, hepatocyte injury, and fibrosis in mouse livers, which suffered NAFLD and NASH under dietary challenge conditions. Orm2 knockout mice spontaneously underwent obese after 16 weeks under normal diet. Orm2 deficiency exacerbated HFD-induced steatosis, steatohepatitis, and fibrosis in mice. Mechanistically, the deletion of Orm2 led to the activation of the Erk1/2-PPARγ-Cd36 signaling pathway, which in turn increased fatty acid uptake and absorption in hepatocytes and mice. Collectively, our results provide Orm2 is an essential factor for preventing NASH and associated NAFLD under obesity.

## Introduction

The number of obese and overweight individuals has increased dramatically over the past decades. Non-alcoholic fatty liver disease (NAFLD) is the most common chronic liver disease globally and pathologically ranges from simple steatosis to non-alcoholic steatohepatitis (NASH) with or without hepatic fibrosis, in the absence of excessive alcohol intake ^1^. While steatosis is not associated with a significant increase in liver-related morbidity or mortality, NASH can progress to more severe diseases, such as cirrhosis and hepatocellular carcinoma, leading to liver failure that requires liver transplantation ^2,3^. It is estimated that approximately 1% of the western population may suffer from NASH. Currently, there are no FDA-approved drugs for the treatment of NAFLD, creating an urgent medical need ^4^.

Increased free fatty acid uptake and de novo lipogenesis account for the majority of the hepatic fat source in NAFLD. Although the mechanisms explaining the association between NAFLD and NASH remain unclear, emerging evidence indicates that NAFLD alters the secretome of liver, including hepatokines, lipids, and non-coding RNAs ^5^. Hepatokines are proteins secreted by hepatocytes, and many hepatokines have been linked to the induction of metabolic dysfunction. The secretion of hepatokines can influence metabolic processes through autocrine, paracrine, and endocrine signaling. These proteins include selenoprotein P, sex hormone-binding globulin (SHBG), fibroblast growth factor 21 (FGF21), and adropin, and so on ^6,7^. Hepatic steatosis induces changes in hepatokine secretion, which promotes insulin resistance and negatively affects other metabolic processes. A better understanding of hepatokine function in hepatic steatosis will aid in the prevention, diagnosis, and treatment of a range of metabolic diseases, including type 2 diabetes mellitus (T2DM). For instance, SHBG is a liver-secreted protein linked to impaired metabolism ^8,9^. SHBG secretion has been inversely associated with liver steatosis, cardiometabolic risk, and insulin resistance, and low levels of SHBG are strong predictors of T2DM risk ^10,11^.

In this study, we accidentally observed that mice with high Oromucoid 2 (Orm2) expression were resistant to high fat diet (HFD)-induced obesity and NAFLD, implying beneficial effects of Orm2 in the liver. Orosomucoid (Orm) is an acute-phase reactive protein and an important secreted glycoprotein molecule synthesized mainly by the liver ^12^. It can be found in plasma under various pathological conditions such as infection, inflammation, tumor, tissue injury, and sepsis. There are two types of Orm, Orm1 and Orm2 ^13^. The Orm2 protein levels are significantly increased in patients with chronic fatigue syndrome (CFS) ^14^. Orm2 inhibits liver cancer cell migration and invasion *in vitro* and intrahepatic metastasis *in vivo* ^15^. Orm2 exerts anti-inflammatory effects by regulating the activation and migration of microglia during brain inflammation, thereby confirming its potential to be used to treat inflammatory diseases ^16^. The above examples highlight that Orm2 functions as a protective factor in different diseases. Given that Orm2 is primarily synthesized in the liver, a more direct effect may exist in the liver through autocrine signaling.

Our study demonstrated that recombinant ORM2 protein inhibited the progression of fatty liver and inflammation, and exhibited significant therapeutic effects in NAFLD and NASH mouse models. The results highlight the critical role of Orm2 protein in the regulation of fatty acid uptake and absorption and provide new insights into the treatment of obesity, NAFLD, and other related metabolic diseases.

## Results

### Mice with high level of Orm2 are resistant to high fat diet-induced obesity

Over the course of 16 weeks of feeding mice a high-fat diet (HFD), we noticed that two mice were resistant to HFD-induced obesity (**Figure 1A**). The two mice (lean group) exhibited weight changes close to those of the chow diet mice (**Figure S1A**). The average daily food intake of the lean group mice was similar to that of the fat group mice under HFD (**Figure S1C**). The serum results showed that lean mice had lower total cholesterol (TC), glucose, triglyceride (TG), HDL cholesterol, and LDL cholesterol levels (**Figure 1B, S1B**). Hematoxylin and eosin (H&E) staining showed less lipid accumulation in the livers of lean mice than in those of fat mice (**Figure 1C**). Furthermore, CD11b immunofluorescence staining showed that inflammatory cell markers were significantly lower in lean mice (**Figure 1D**). To further systematically investigate how lean mice are protected from HFD-induced fatty liver, we performed RNA-Seq on the livers of mice. A heatmap revealed that the expression of hepatic genes related to inflammation and lipid metabolism pathways was significantly downregulated in lean mice with HFD for 16 weeks (**Figure 1E**). After screening the genes with a high fold difference and expression between fat and lean mice (**Figure 1F**), we focused on Orm2, which was elevated 52-fold between fat and lean mouse livers. Changes in Fasn, Atgl, and Orm2 expression were further confirmed by western blotting, which showed that lean mice had higher fatty acid hydrolysis and lower fatty acid synthesis (**Figure 1G**).

**Figure 1.**
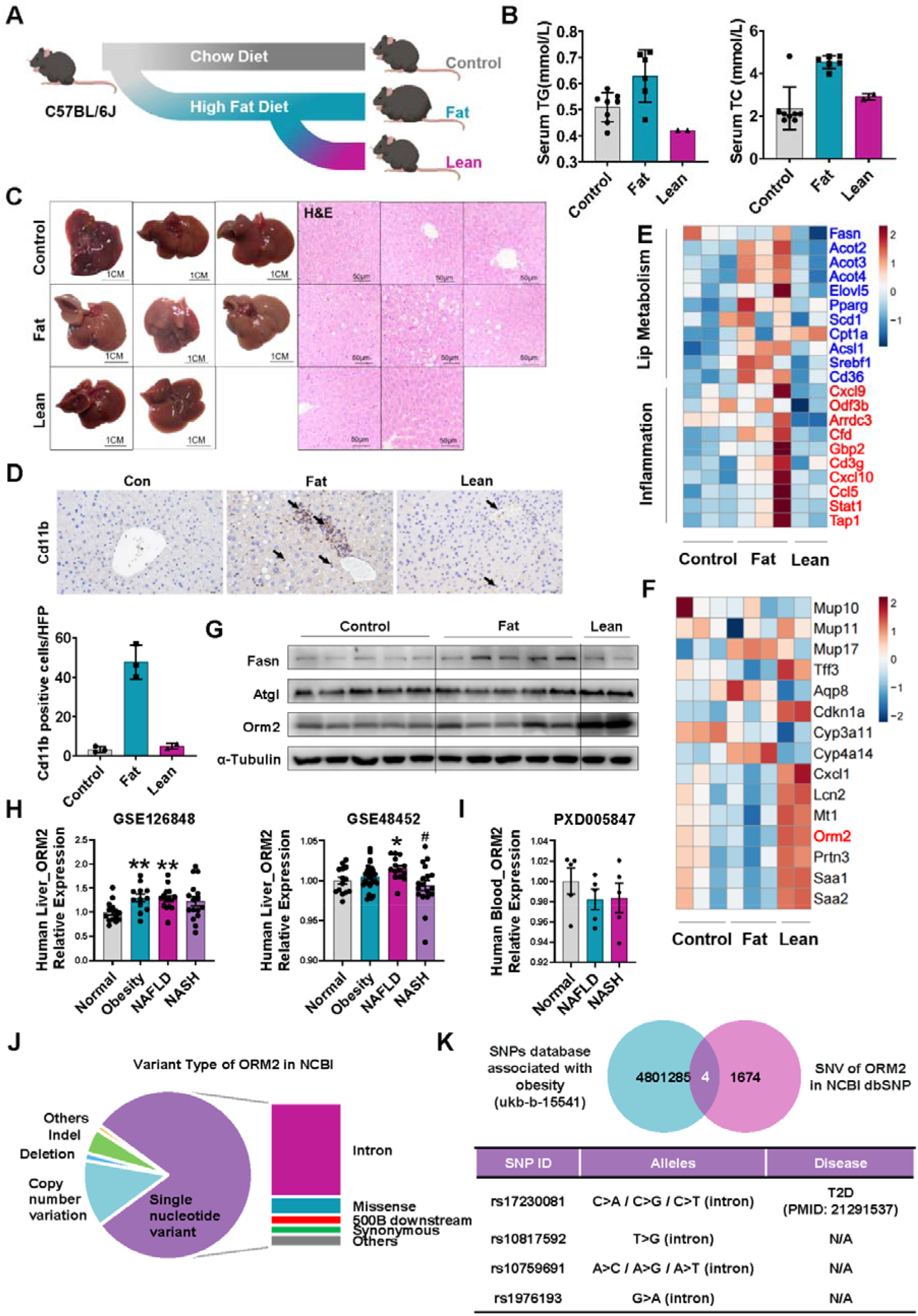
Mice with high level of Orm2 are resistant to high fat diet-induced obesity. (A) Schematic diagram of mice with or without high-fat diet. (B) Serum cholesterol and triglycerides levels. (C) Representative pictures of liver and images of H&E stained sections. Scale bar 100 μm. (D) Representative immunohistochemical image of Cd11b, and quantitative results by Image-J (n = 2-3/group). (E) Heatmap of the liver lip metabolism and inflammation genes detected by RNA-seq. (F) Heatmap of differential genes. (The FPKM value is greater than 5, and the fold difference between HFD and Lean is greater than 10). (G) Western blotting of Fasn, Atgl, and Orm2 in liver. Protein band density was determined by Image J software and normalized to ɑ-Tubulin (bottom). (n = 2 for Lean; n = 3 for the other groups). (H) Analysis of ORM2 expression in the human obesity-related liver RNA-seq dataset from the GEO database (https://www.ncbi.nlm.nih.gov/gds/?term=). (I) The serum ORM2 levels in the PXD005847 dataset in the ProteomeXchange database (http://proteomecentral.proteomexchange.org/cgi/GetDataset). (J) Mutations in ORM2 in the NCBI database (https://www.ncbi.nlm.nih.gov/variation/view/). (K) Combine the obesity-related SNP dataset (ukb-b-15541) in the Open GWAS database (https://gwas.mrcieu.ac.uk/) with the NCBI dbSNP database (https://www.ncbi.nlm.nih.gov/snp/?term=) for ORM2 SNV data for intersection analysis. All analyzed data were normalized to the Normal group. *indicates a significant difference between the Normal group and the Non-Normal group; \**P* < 0.05, \*\**P* < 0.01. # indicates a significant difference between the NAFLD group and the NASH group; #*P* < 0.05.

In the NCBI database, Orm2 expression was significantly upregulated in the livers of obese patients (**Figure 1H, 1I, and Figure S2A**). Orm2 expression declined as NAFLD progressed to NASH, whereas it decreased after bariatric surgery and control of the caloric intake (**Figure S2B**). The expression of Orm2 was also upregulated in obese mice, including db/db, ob/ob, and HFD-induced obese mice (**Figure S2C**), further suggesting that Orm2 is a potential target for obesity. Moreover, a combined analysis of Orm2 potential SNP sites and obesity-related SNP datasets revealed four intron mutations, of which the rs17230081 site was a potential regulatory site affecting the development of T2DM (**Figure 1J, 1K**) ^17^.

The sequencing of Orm2 gene promoter showed that the promoter region of Orm2 was not mutated in the lean mice (**Figure S3A**). STAT1 has been confirmed by multiple studies and is significantly upregulated in the NASH ^18^. The analysis of NASH patient data (GSE173734_ATAC-seq from NCBI) showed that the STAT1-binding region of the Orm2 promoter showed an open chromatin status (**Figure S3B**). We further verified that STAT1 can bind to the Orm2 promoter region to transactivate transcription using a dual-luciferase reporter gene assay (**Figure S3C, S3D**). In addition, the upregulation or downregulation of STAT1 altered the expression of Orm2 correspondingly (**Figure S3E, 3F**). These results suggested that STAT1 may be a key factor in the upregulation of Orm2 in hepatocytes.

### ORM2 protein treatment ameliorated HFD-induced NAFLD and metabolic dysfunction in mice

The extremely high Orm2 expression in lean mouse livers implies that Orm2 may have beneficial effects on HFD-induced NAFLD. To gain insight into the therapeutic role of Orm2 in the liver, long-term usage of recombinant ORM2 protein was performed in HFD mice. The ORM2 protein treatment significantly reduced the body weight, liver weight, and adipose weight in mice (**Figure 2A-D**). Decreased contiguous patches of microsteatosis, hepatocyte ballooning, and accumulation of hepatic lipids were revealed by H&E and Oil Red O staining in the ORM2-treated HFD mice (**Figure 2E**). Notably, ORM2 reduced hepatic steatosis, the lipid content, and the acute plasma TC, TG, AST, and ALT levels (**Figure 2F**). Furthermore, the HFD-induced increase in lipid-related gene expression was abrogated by the ORM2 treatment (**Figure 2G**). ORM2-treated mice had improved glucose tolerance and insulin sensitivity, indicating improvement in metabolic dysfunction (**Figure 2H and 2I**). These results indicate that ORM2 protects the mice from the pathological damage that leads to the development of NAFLD.

**Figure 2.**
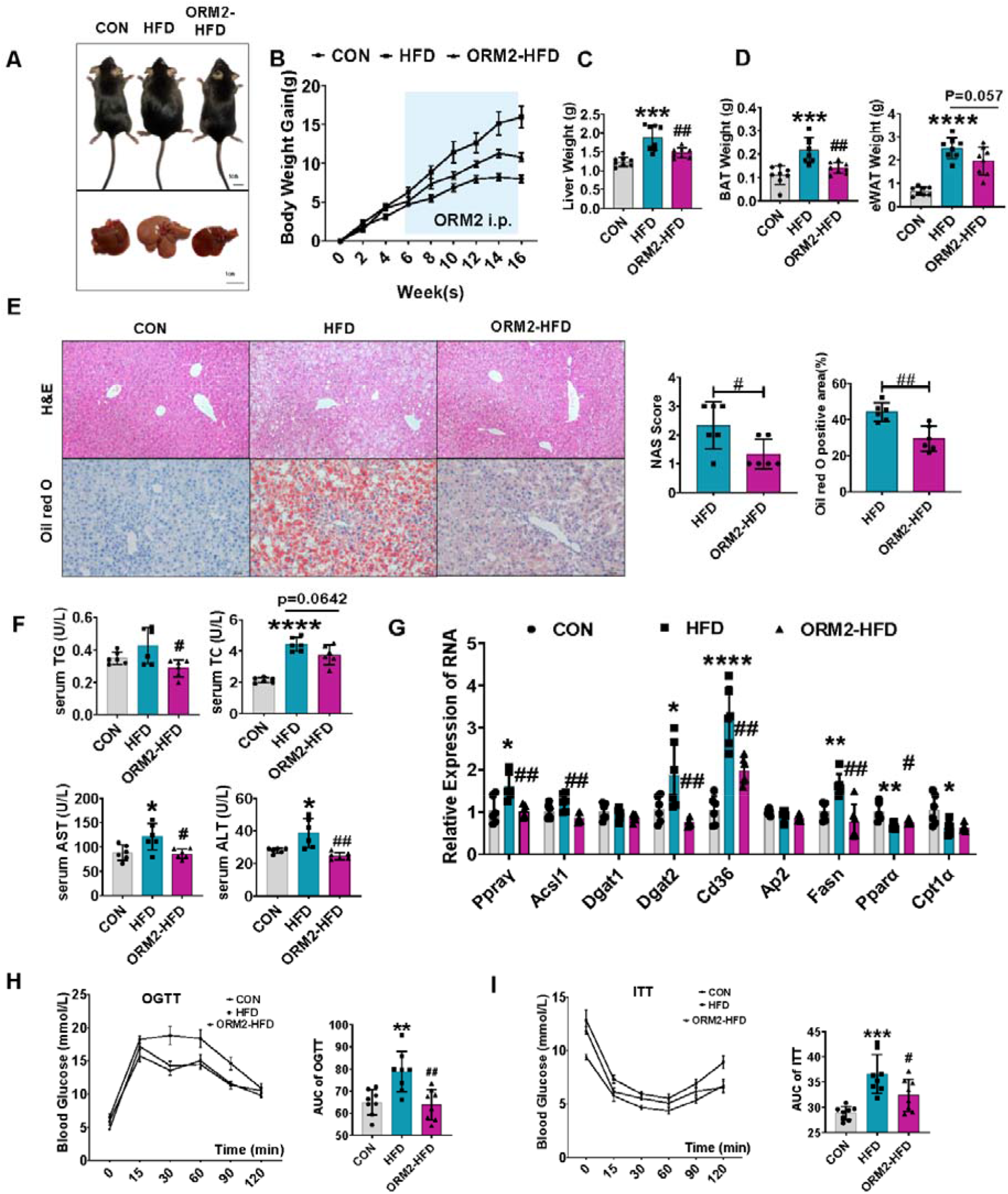
Orm2 protein treatment ameliorates HFD-induced NAFLD and metabolic dysfunction in mice. (A) Representative appearance pictures of mice (top), liver (bottom). (B) Body weight change in CON (fed normal chow diet), HFD, and ORM2-HFD (fed high-fat diet), n = 8 per group. Liver weight, (D) BAT and eWAT in mice at 24 weeks of age, n =6 per mice group. (E) Representative H&E and Oil red O staining and quantified results, (n = 6/group). Scale bar 100 μm for H&E staining and 50 μm for Oil red O staining. (F) Serum ALT, AST, TC, and TG levels were measured in mice at 24 weeks. n =6 per group. (G) RT-qPCR for the indicated metabolism genes expression organized for liver. Data are expressed as fold-change in expression relative to CON following calculation of relative expression to Gapdh. (H) OGTT ITT and area under curve (AUC) shown. n = 8 per group. Data are the mean±S.E.M. and were compared by Student’s t-test. *indicates a significant difference between the CON group and the HFD group; \**P* < 0.05, \*\**P* < 0.01, \*\*\**P* < 0.001, and \*\*\*\**P* < 0.0001. # indicates a significant difference between the ORM2-HFD group and the HFD group; #*P* < 0.05, ##*P* < 0.01.

### ORM2 protein treatment ameliorated MCD-induced NASH in mice

Given the function of Orm2 in HFD-induced NAFLD, we investigated the role of Orm2 in the development of steatohepatitis. The recombinant ORM2 protein treatment was implemented in MCD diet induced hepatitis mice. Compared with control mice, mice that were given a MCD diet for four weeks had enlarged and yellowish livers, a dramatic loss of body weight, and an increased liver-to-body weight ratio, which was reversed by the administration of ORM2 (**Figure 3A-3C**). H&E staining, Sirius Red staining, and Oil Red O staining showed that Orm2-treated mice had significantly reduced liver steatosis, necrotic inflammation, and deposition of collagen fibers compared with untreated mice with MCD diet (**Figure 3D**). The accumulation of TC and TG in the liver was significantly reduced after the ORM2 treatment (**Figure 3E**). The serum AST and ALT levels also showed the ORM2 resistance to MCD-induced liver injury (**Figure 3F**). In addition, Cd11b staining revealed that ORM2 treatment attenuated MCD-induced liver inflammation (**Figure 3G**). RT-qPCR assays also confirmed that Orm2 inhibited the expression of pro-inflammatory cytokines (Mcp1, Cxcl10, and IL1β) induced by the MCD diet (**Figure 3H**), and western blotting detected decreased fibrosis (Collagen 1 and ◻-SMA) (**Figure 3I**). These results indicate that ORM2 treatment ameliorated NASH in mice.

**Figure 3.**
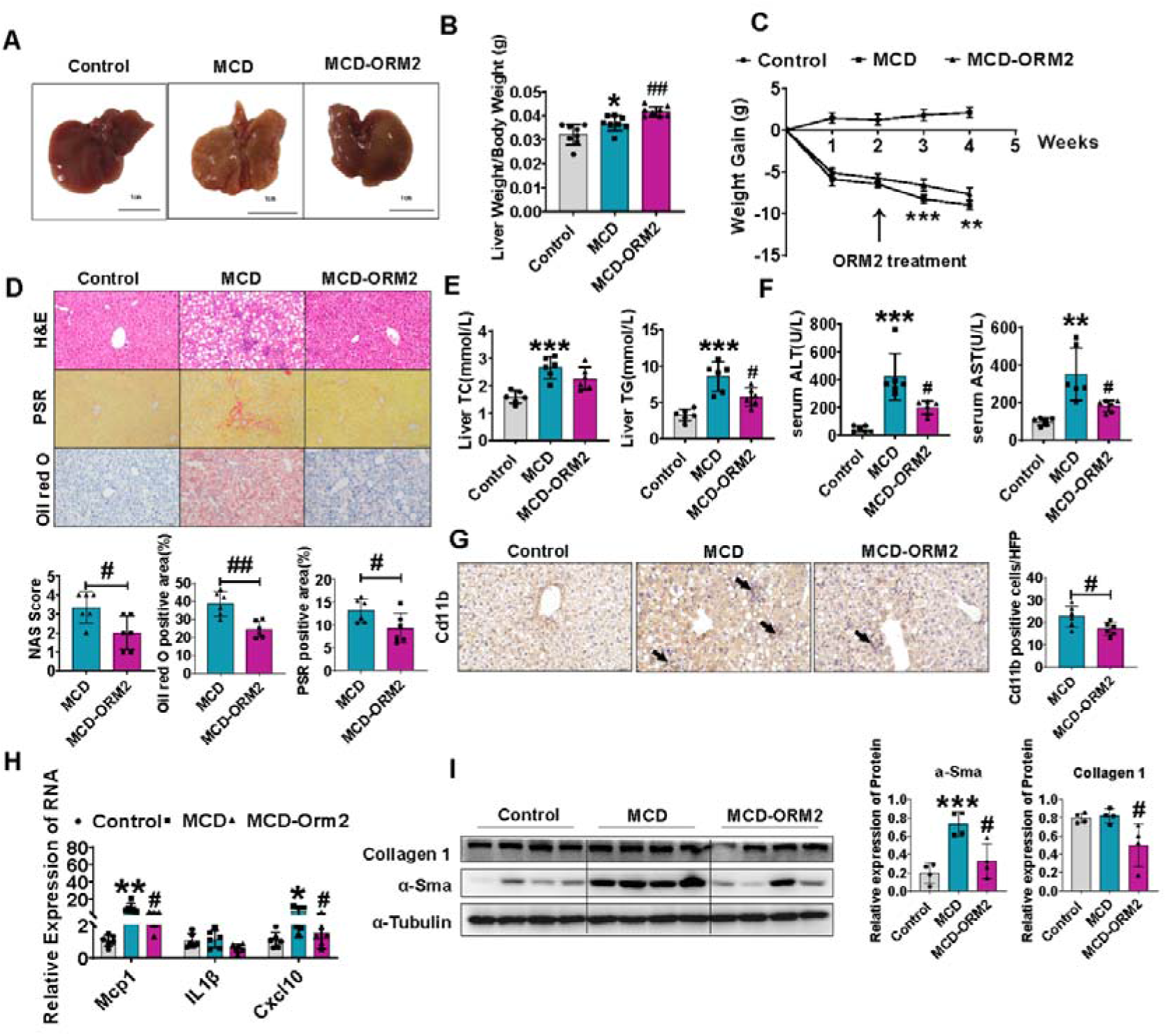
ORM2 protein treatment ameliorated MCD-induced NASH in mice. (A) Representative appearance pictures of liver in Control (fed regular chow), MCD, and MCD-ORM2 (fed methionine and choline deficient L-amino acid diet). (B) Liver Weight / Body Weight in mice at 12 weeks of age among Control, MCD, and MCD-ORM2, n = 8 per group. (C) Body weight change in Control, MCD, and MCD-ORM2. (D) Representative images of H&E staining (top), Picrosirius Red staining (middle) and Oil red O staining (bottom) of liver and quantitative results from Image-J (n = 6/group). Scale bar 50 μm for Oil red O staining, Scale bar 100 μm for the others. (E) Hepatic lipid (TG and TC) levels among Control, MCD, and MCD-ORM2. n =8 per group. (F) Serum ALT and AST levels were measured in mice at 12 weeks. n =6 per group. (G) Representative immunohistochemical image of Cd11b, and quantitative results from Image-J (n = 6/group). (H) Quantitative PCR for the indicated inflammation genes organized for liver. Data are expressed as fold-change in expression relative to Control following calculation of relative expression to Gapdh. n =6 per group. (I) Western blotting analysis of the Collagen 1 and α-Sma protein for three groups. α-Tubulin was used as internal control. Data are the mean±S.E.M. and were compared by Student’s t-test. * indicates a significant difference between the Control group and the MCD group; \**P* < 0.05, \*\**P* < 0.01, \*\*\**P* < 0.001. # indicates a significant difference between the ORM2-MCD group and the MCD group; #*P* < 0.05, ##*P* < 0.01.

### Orm2 KO caused spontaneous weight gain and metabolic dysfunction in mice

Orm2 is mainly expressed in the liver and rarely expressed in other tissues. To directly address the function of Orm2 in the liver, we generated Orm2 KO mice. As representative images of mouse epididymal fat and liver showed in **Figure 4A**, Orm2 KO mice had higher body weight and epididymal white adipose tissue (eWAT) and brown adipose tissue (BAT) weights compared to wild type mice (WT) at 20 weeks (**Figure 4B, 4C**). H&E staining confirmed the enlargement of adipocytes (**Figure S4A**). However, no obvious changes were observed in liver weights (**Figure 4D**), fat deposition in hepatocytes (**Figure 4E**), fat deposit or metabolism genes (**Figure 4F**). No significant inflammatory cell infiltration in hepatic tissue was observed either (**Figure S4B**). Consistent with the hepatic pathological results, there is no significant alteration in the expressions of fatty acid synthesis (PPARγ, Acsl1, Dgat1, Dgat2, and Fasn), fatty acid uptake (Cd36 and Ap2), and fatty acid hydrolysis (Cpt1◻, PPAR◻) genes (**Figure 4F**). Next, we evaluated the metabolic profile of Orm2 KO mice by oral glucose tolerance and intraperitoneal insulin-tolerance tests (O-GTT and IP-ITT, respectively). Orm2 KO mice exhibited impaired insulin sensitivity but not glucose tolerance (**Figure 4G and Figure S4C**). Furthermore, the average daily food intake of the Orm2 KO mice did not change for four weeks (**Figure S4D**). These results suggest that Orm2 KO mice had spontaneous obesity and metabolic dysfunction but did not exhibit profound hepatic steatosis in a standard diet.

**Figure 4.**
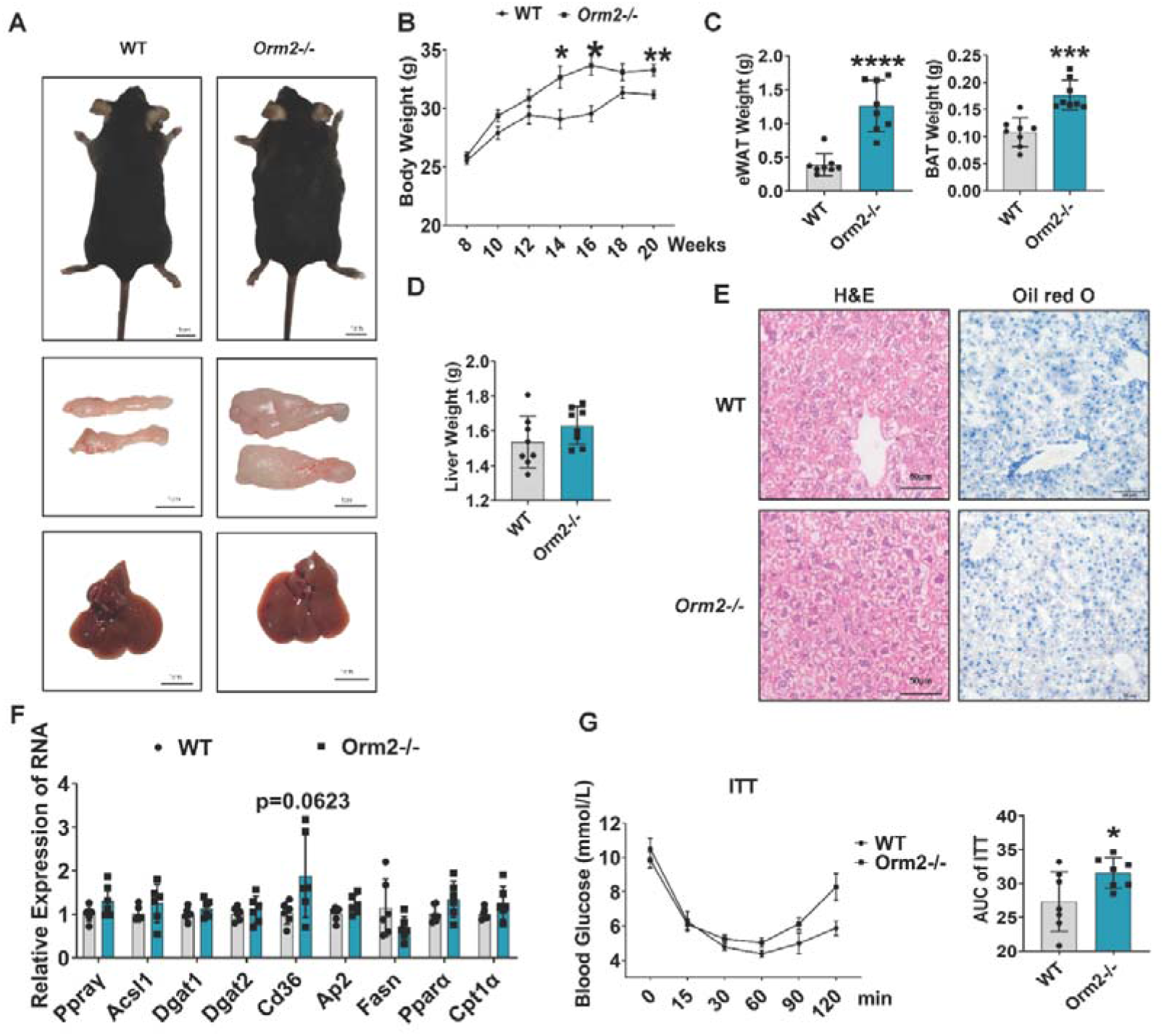
Orm2 KO caused spontaneous weight gain and metabolic dysfunction in mice. (A) Representative appearance pictures of mice (top), epididymal white fat (middle), and liver (bottom). (B) Body weight change in WT and Orm2 KO mice during regular chow. (C, D) eWAT weight, BAT weight, and liver weight of WT and Orm2 KO mice during a regular chow diet for 20 weeks (n = 8/group). (E) Representative images of H&E (left) and Oil red O (right) staining of liver. Scale bar 50 μm. (F) RT-qPCR for the indicated metabolism genes organized for liver. Data are expressed as fold-change in expression relative to WT following calculation of relative expression to Gapdh. (G) Intraperitoneal insulin tolerance test (ITT) performed at 18 weeks of age; glucose measurements over time after insulin bolus-injected i.p. (left) and AUC (right) shown. n = 8 per group. All data are shown as mean±S.E.M. and were compared by Student’s t-test, where \**P* < 0.05, \*\**P* < 0.01, \*\*\**P* < 0.001, and \*\*\*\**P* < 0.0001.

### Orm2 KO mice exacerbated diet-induced NAFLD and NASH

Given that the liver health of Orm2 KO mice was not affected on a standard diet, we detected the effects of Orm2 KO on the liver by changing the dietary conditions. Male WT and Orm2 KO mice were fed the HFD for 16 weeks. Compared with WT mice, Orm2 KO mice showed higher body weight (**Figure 5A, 5B**) and exhibited an increased liver weight to body weight ratio and adipose weight (**Figure S5A**), as well as an increase in adipocyte volume (**Figure S5B**). Orm2 KO mice fed with HFD for 16 weeks exhibited significantly higher hepatic total cholesterol (TC) and triglyceride (TG) levels (**Figure 5C**). In addition, contiguous patches of micro-steatosis, hepatocyte ballooning, lobular inflammation, and accumulation of hepatic lipids were revealed by H&E and Oil Red O staining in Orm2 KO mice (**Figure 5D**). However, immunofluorescence staining revealed no obvious high CD11b (+) cell infiltration (**Figure S5C**). Next, we examined the hepatic expressed genes that related to fatty acid synthesis (PPARγ, Dgat2, and Fasn) and fatty acid uptake (Cd36 and Ap2), which elevated in Orm2 KO mice than in control mice after 16 weeks of HFD feeding (**Figure 5E**). Serum TG, TC, ALT, and AST levels were higher in Orm2 KO mice than in WT mice, which implied the obesity and hepatic damage (**Figure 5F**). Additionally, insulin sensitivity was further evaluated using ITT, which showed that Orm2 KO mice had decreased insulin sensitivity (**Figure 5G**). These results indicated that Orm2 KO exacerbated HFD-induced obesity and NAFLD.

**Figure 5.**
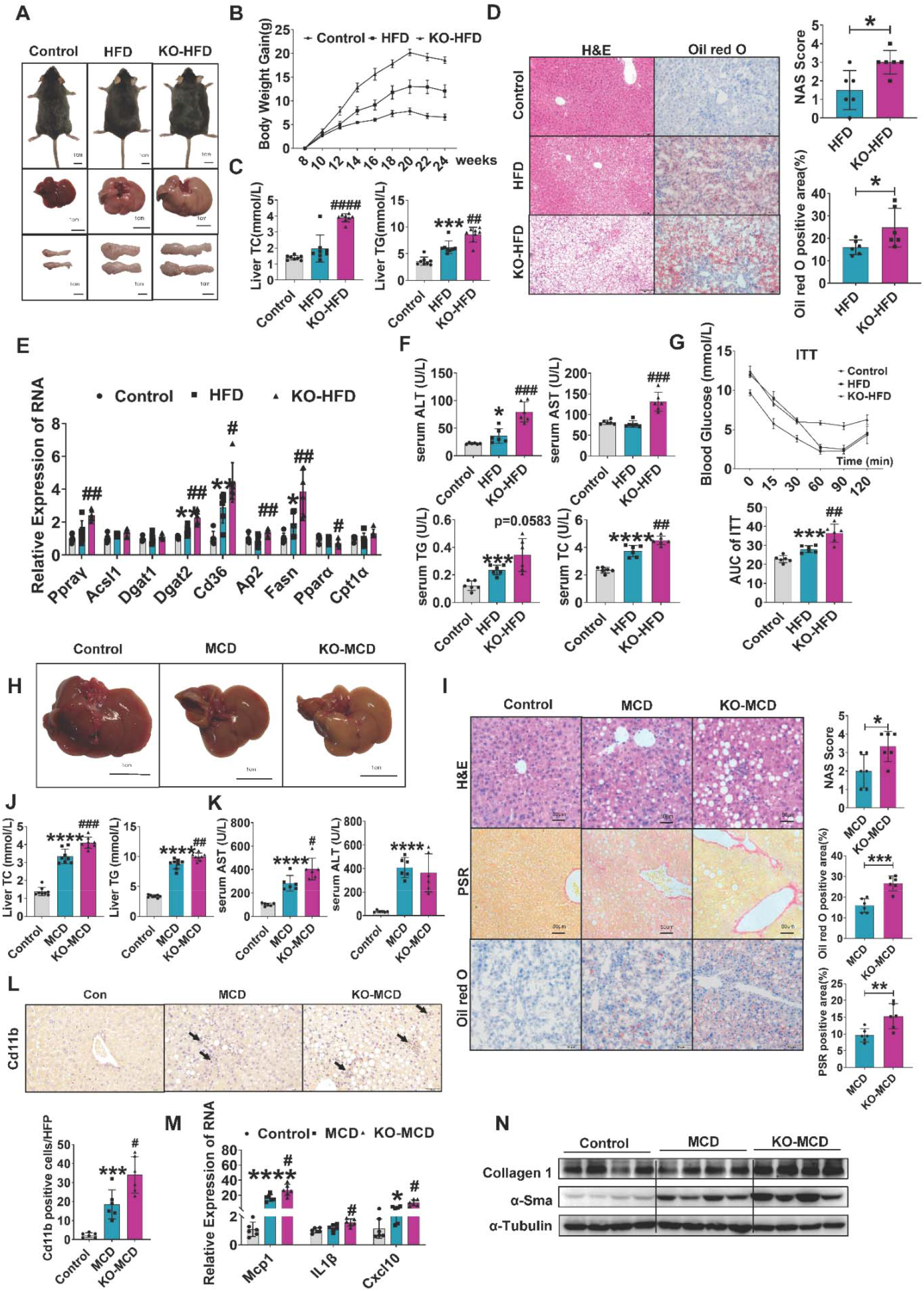
Orm2 KO mice exacerbated diet-induced NAFLD and NASH. (A) Representative appearance pictures of mice (top), liver (middle), and epididymal white fat (bottom). (B) Body weight change in Control (fed regular chow), HFD, and KO-HFD (fed high-fat diet), n = 8 per group. (C) Hepatic lipid (TG and TC) levels. n =8 per group. (D) Representative H&E and Oil red O staining and quantified results of Control (fed regular chow), HFD, and KO-HFD (fed high-fat diet) (n = 6/group). (E) RT-qPCR for the indicated metabolism genes organized for liver. Data are expressed as fold-change in expression relative to Control following calculation of relative expression to Gapdh. (F) Serum ALT, AST, TC, and TG levels were measured in mice at 24 weeks. n =6 per group. (G) ITT performed at 22 weeks of age; glucose measurements over time after insulin bolus-injected i.p. (left) and AUC (right) shown. Data are the mean±S.E.M. for n = 6 mice per group and were compared by Student’s t-test. (H) Representative appearance pictures of liver in Control (fed regular chow), MCD, and KO-MCD (fed methionine and choline deficient L-amino acid diet). (I) Representative images of H&E staining (top), Picrosirius Red staining (middle) and Oil red O staining (bottom) of liver and quantitative results from Image-J (n = 6/group). Scale bar 50 μm. (J) Hepatic lipid (TG and TC) levels among Control, MCD, and KO-MCD. (K) Serum ALT and AST levels were measured in mice at 12 weeks. n =6 per group. (L) Representative immunohistochemical image of Cd11b, and quantitative results from Image-J (n = 6/group). (M) RT-qPCR for the indicated inflammation genes organized for liver. Data are expressed as fold-change in expression relative to Control following calculation of relative expression to Gapdh. n =6 per group. (N) Western blotting analysis of the Collagen 1 and α-Sma protein levels for three groups. α-Tubulin was used as internal control. Data are the mean±S.E.M. and were compared by Student’s t-test. * indicates a significant difference between the Control group and the HFD/MCD group; \**P* < 0.05, \*\**P* < 0.01, \*\*\**P* < 0.001, \*\*\*\**P* < 0.0001. # indicates a significant difference between the KO-HFD/MCD group and the HFD/MCD group; #*P* < 0.05, ##*P* < 0.01, ###*P* < 0.001, ####*P* < 0.0001.

Next, MCD diet was used to address the role of Orm2 in the progression of NAFLD to NASH. Orm2 KO and WT mice were fed a control diet or a MCD diet for four weeks to trigger steatohepatitis. The MCD diet resulted in enlarged and yellowed livers, increased liver/weight ratio, and significant weight loss in mice. This phenomenon was exacerbated in Orm2 KO mice (**Figure 5H, Figure S5D, and Figure S5E**). Consistently, H&E and Oil Red O staining showed a significant increase in hepatic steatosis and necroinflammation in Orm2 KO mice compared with WT mice fed MCD (**Figure 5I**), which was also supported by the accumulation of TC and TG in the liver (**Figure 5J**). In contrast to WT mice, knockout of Orm2 expression increased the serum levels of AST and ALT (**Figure 5K**). Additionally, Cd11b and PSR staining revealed that Orm2 KO exacerbated MCD-induced liver inflammation and collagen deposition (**Figure 5L**). RT-qPCR assays also confirmed that Orm2 KO increased the mRNA levels of inflammation-related genes (**Figure 5M**), and western blotting detected elevated fibrosis (**Figure 5N**). These results suggested that Orm2 deficiency aggravates NASH significantly.

### Orm2 deletion activated the Erk-PPARγ-Cd36 signaling pathway in the mouse liver

To further explore the mechanism by which Orm2 deficiency exacerbates HFD-induced fatty liver, we analyzed the liver transcriptomes of Orm2 KO mice and WT mice treated with HFD for 16 weeks. Heatmap showing genes of lipid metabolic process enriched in GO term are most highly expressed in Orm2 KO mice (**Figure 6A**). Subsequently, KEGG was used to analyze the enriched signaling pathways and the scores of PPARγ signaling pathway ranked in the top three pathways (**Figure 6B**). Further analysis showed that genes involved fatty acid uptake and synthesis in the PPARγ signaling pathway were higher in the livers of Orm2 KO mice than in those of the other two groups (**Figure 6C**).

**Figure 6.**
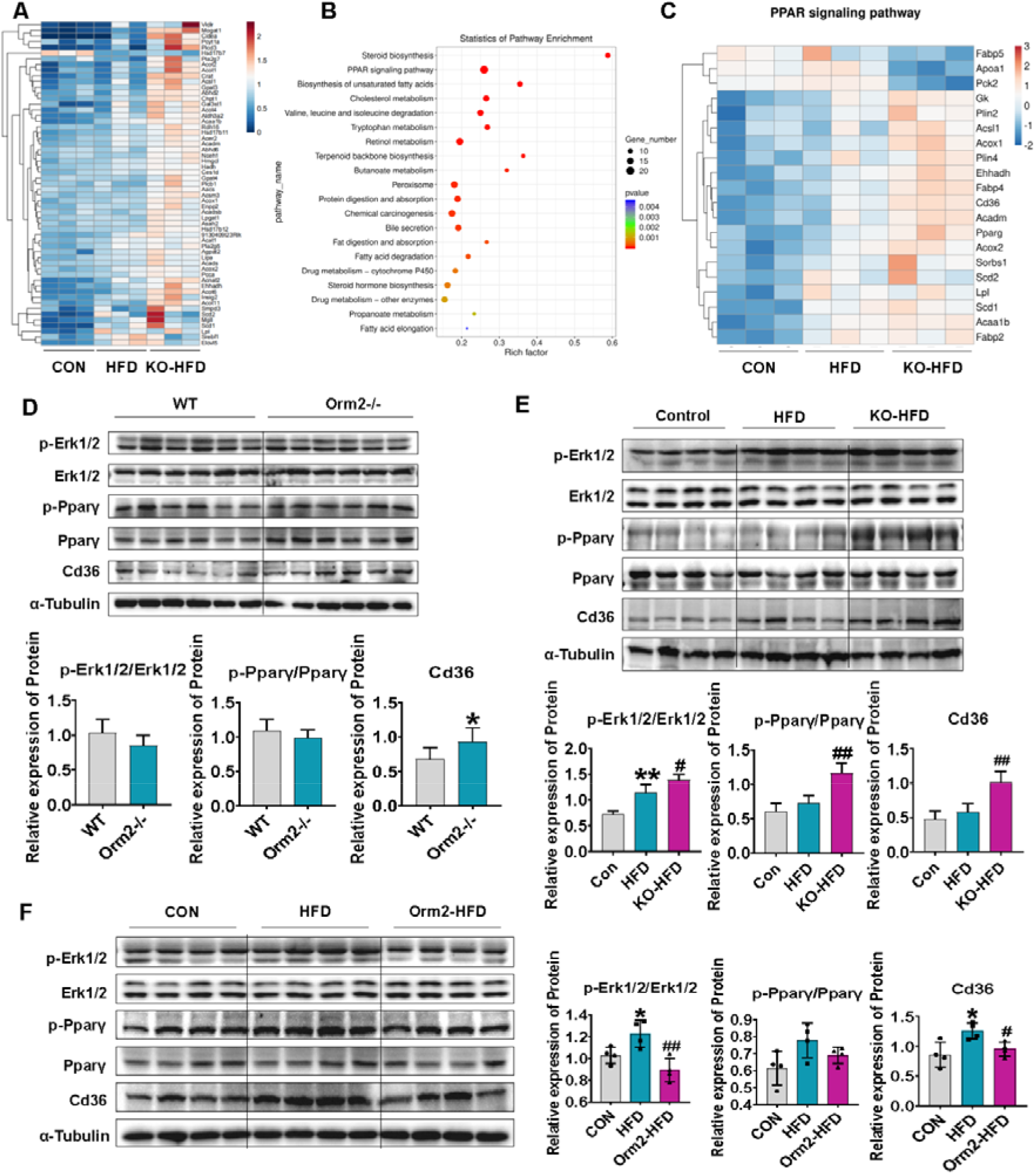
Orm2 deletion activated the Erk-PPARγ-Cd36 signaling pathway in the mouse liver. (A) Heatmap showing lipid metabolic process enriched in GO term in RNA-seq from liver. n = 3 per group. (B) Bubble plot showing three groups’ KEGG pathway enrichment in RNA-seq. (C) Heatmap showing changes in genes enriched in PPAR signaling pathway in RNA-seq. (D) Representative western blotting of p-Erk1/2, Erk1/2, p-PPARγ, PPARγ, and Cd36 in liver between WT and Orm2 KO mice (top). α-Tubulin was used as internal control. Protein band density was determined by Image J software and normalized to α-Tubulin (bottom). n = 6 per group. Data are the mean±S.E.M. (E) Representative western blotting of p-Erk1/2, Erk1/2, p-PPARγ, PPARγ, and Cd36 in liver among Control, HFD, and KO-HFD mice. (F) Western blotting analysis of the Cd36, total and phosphorylated protein levels of PPARγ and Erk1/2 among CON, HFD and ORM2-HFD. Protein band density was determined by Image J software and normalized to α-Tubulin (bottom). n = 4 per group. Data are the mean±S.E.M. and were compared by Student’s t-test. * indicates a significant difference between the WT group and the ORM2-/-group, or the Control group and the HFD group; \**P* < 0.05, \*\**P* < 0.01. # indicates a significant difference between the KO-HFD group and the HFD group, or the ORM2-HFD group and the HFD group; #*P* < 0.05, ##*P* < 0.01.

Many studies have confirmed that PPARγ regulates the expression of Cd36 and plays an active role in the occurrence of fatty liver ^19,20^. Moreover, Erk1/2 plays an important role in NAFLD and induces hepatic lipid accumulation ^21^. First, the protein expression of Erk1/2, PPARγ, and Cd36 was examined in the livers of mice fed a normal diet. The results showed that Orm2 KO did not affect the expression of Erk1/2 and PPARγ in the liver but increased the expression of Cd36 (**Figure 6D**). However, under the high-fat diet, Erk1/2, PPARγ, and Cd36 expression increased in the liver of Orm2 KO mice (**Figure 6E**). The elevation of p-Erk1/2, p-PPARγ, and Cd36 induced by HFD was attenuated by the recombinant ORM2 treatment (**Figure 6F**). These results suggest that Orm2 KO activated the Erk1/2-PPARγ-Cd36 signaling pathway to promote fatty acid uptake and absorption under HFD conditions, but not under the standard chow diet.

### Orm2 inhibited fatty acid uptake by inhibiting Erk1/2-PPARγ-Cd36 in hepatocytes

To further assess the function of Orm2 in fatty liver, *in vitro* experiments were performed using human and mouse hepatocytic cell lines. First, Orm2 small-interfering (si)RNA or Orm2 overexpression plasmid was transfected in human LO2 hepatocytes. Revealed by Oil Red O staining, Orm2 silencing increased OA/PA-induced lipid storage, and Orm2 overexpression decreased the lipid content in LO2 cells (**Figure 7A**). Similarly, the interference or overexpression of Orm2 significantly altered the OA/PA-induced lipid-related gene expression (**Figure 7B**). To investigate whether Orm2 mediates the lipid accumulation regulation of PPARγ activation, we tested the effect of the PPARγ agonist pioglitazone (Pio) on the phosphorylation of PPARγ and its downstream target Cd36. Erk1/2, PPARγ, and Cd36 were evaluated in cultured AML12 mouse hepatocytes exposed to OA/PA and recombinant mouse ORM2 protein. The co-treatment with Orm2 and OA/PA significantly decreased the OA/PA-induced activation of the Erk1/2-PPARγ-Cd36 signaling pathway and abolished the effect of Orm2 on Cd36 and PPARγ inhibition but did not affect the Erk1/2 (**Figure 7C**). Lipid metabolic genes in Orm2-treated hepatocytes in the presence or absence of Pio were examined by RT-qPCR assays. The downregulation of fatty acid synthesis and uptake genes (Cd36, PPARγ, Acsl1, and Fasn) by Orm2 was reversed by Pio (**Figure 7D**). These results suggested that Orm2 regulates hepatocyte lipid uptake through an Erk1/2-PPARγ-Cd36 signaling pathway.

**Figure 7.**
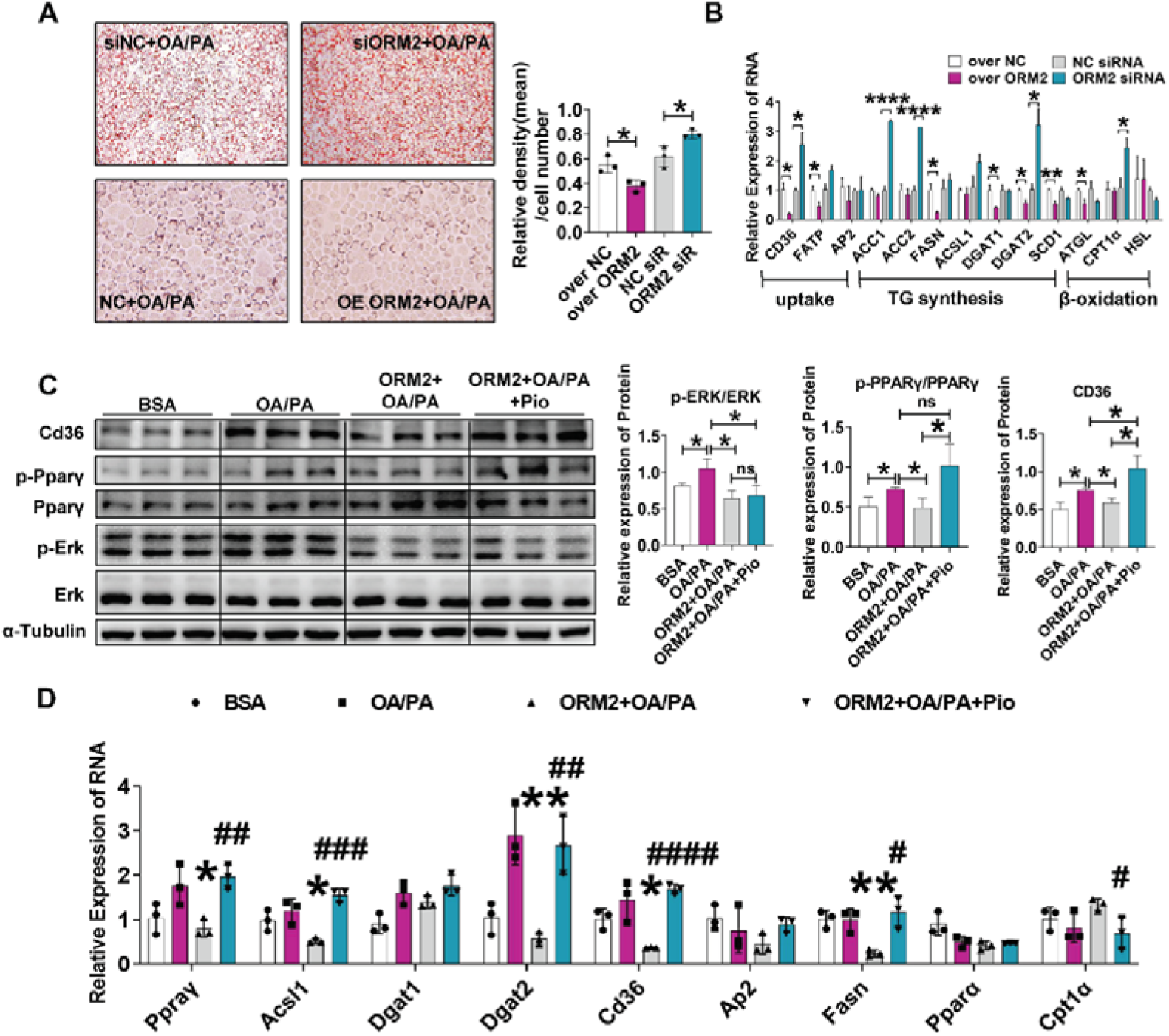
Orm2 inhibited fatty acid uptake by inhibiting Erk1/2-PPARγ-Cd36 in hepatocytes. (A) Representative Oil red O staining of LO2 cells with ORM2 downregulated or ORM2 overexpressed, treated with OA/PA mixture (OA 0.4 mM, PA 0.2 mM) for 24 h (n = 3 per group). (B) RT-qPCR for the indicated metabolism genes organized for LO2 cells with ORM2 downregulated or ORM2 overexpressed, treated with OA/PA mixture (OA 0.4mM, PA 0.2mM) for 24 h (n = 3 per group). Data are expressed as fold-change in expression relative to Control following calculation of relative expression to Gapdh. (C) Western blotting analysis of the Cd36, total and phosphorylated protein levels of PPARγ and Erk1/2 after treatment with OA/PA (OA 0.4 mM, PA 0.2 mM) for 12 h with or without the addition of ORM2 protein (500 ng/mL) and PPARγ agonist pioglitazone for AML12 cells. α-Tubulin was used as internal control. n = 3 per group. (D) RT-qPCR for the indicated metabolism genes organized for AML12 cells. Data are expressed as fold-change in expression relative to Control following calculation of relative expression to Gapdh. Data are the mean±S.E.M. for n = 3 per group and were compared by Student’s t-test. * indicates a significant difference between the BSA group and the OA/PA group, or the two groups identified by the line segment in the figure; \**P* < 0.05, \*\**P* < 0.01, \*\*\*\**P* < 0.0001. # indicates a significant difference between the ORM2+OA/PA group and the OA/PA group; #*P* < 0.05, ##*P* < 0.01, ####*P* < 0.0001.

## Discussion

In this study, we demonstrated the therapeutic value of Orm2 in the treatment of experimental NAFLD and NASH, and revealed the underlying mechanism involving in fatty acid metabolism. Mice with high expression of Orm2 in hepatocytes showed the resistant to HFD-induced steatosis. Recombinant ORM2 protein prevented HFD-induced steatosis and MCD-induced liver damage and fibrosis in mice. Under normal dietary conditions, hepatic Orm2 KO mice appeared to have little effect on the liver, but impaired systemic metabolism. However, under HFD or MCD conditions, Orm2 deficiency resulted in significant aggravation of the disease (liver and whole body). Mechanistically, we found that the detrimental effects of Orm2 deficiency on the liver were mediated by activating Cd36 to increase fatty acid uptake and synthesis through the Erk and PPARγ pathway.

The liver affects glucose and lipid metabolism partly by secreting cytokines, which are termed as hepatokines. NAFLD, in particular, is associated with the altered production of hepatokines ^22^. Hepatokines are important contributors to metabolic diseases and may enable the pathophysiological roles of the liver to be distinguished from those of other tissues. An increased understanding of the roles of the liver and hepatokines in metabolic diseases could lead to the development of improved targeting strategies for their prevention and treatment.

Biological therapies with recombinant proteins are among the most promising treatment for various diseases today ^23^. For liver diseases, many of the pharmacological recombinant protein applications involve hepatokines. For example, as a hepatokine, FGF4 recombinant protein protects liver from NAFLD via the AMP-activated protein kinase-Caspase 6 signaling axis ^24^; The FGF19 and FGF21 administration prevents the pathological progression of NAFLD, including hepatosteatosis and NASH associated with obesity and diabetes ^25,26^. In addition, recombinant proteins have also been widely used in other diseases. For example, recombinant IL-35 protein shows significant therapeutic effect on acute colitis and psoriasis mice ^27^; recombinant ORM2 protein displays inhibitory effect on neuroinflammation ^16^; recombinant BTI protein can be used in the prevention of obesity and hyperglycemia in lipidemia ^28^. The above examples highlight that recombinant proteins have undoubtedly been revealed as effective disease therapies. The results of our biochemical and morphological studies of ORM2 recombinant protein administration in mice indicated that Orm2 inhibits Cd36 expression and lipid absorption, which then showed obvious therapeutic effects on NAFLD and NASH.

There are two isoforms of Orm in humans (Orm1 and Orm2), three isoforms in mice (Orm1, Orm2, and Orm3), and one in rats ^29^. The expression of Orm is mainly regulated by a variety of regulators such as glucocorticoids, interleukin-1, tumor necrosis factor-◻, and interleukin-6 ^30^. The altered expression of Orm2 in NAFLD or obesity is still disputable. In a recent study, Zhou. et al described decreased expression of Orm2 in mice and NAFLD patients ^31^. In this study, the changes in Orm2 were more carefully confirmed. By examining GEO databases, we found that, compared to controls, ORM2 expression was elevated in obese and NAFLD patients. Consistent with the results of Zhou. et al, the expression of Orm2 decreased as NAFLD progressed from simple steatosis to inflammatory NASH. But notably, patients who accepted bariatric surgery or caloric restriction had lower Orm2 expression levels. Orm2 is generally considered an anti-inflammatory and immunomodulatory factor owing to its anti-neutrophil and anti-complement activities ^13,30^. We believe that the relatively elevated Orm2 may be a reflection of the low-grade state of inflammation in NAFLD or obesity, while high Orm2 levels help activate immune cells to clear inflammation. Therefore, the altered expression of Orm2 in NAFLD or NASH requires further study, such as large-scale population-based investigation.

In this study, RNA-seq analysis showed that Cd36 expression in the liver of Orm2 KO mice with NAFLD was significantly upregulated under HFD conditions. Studies have shown that the fatty acid translocase Cd36 is an important mediator of lipid uptake in many tissues ^20,32,33^, and Erk of hepatic steatosis by regulating the FFA uptake rate ^33^. This also explains our results that Orm2 deletion causes the upregulation of Cd36 in hepatocytes and then induces hepatic steatosis and obesity under the HFD. Previous studies reported that PPARγ synergistically promotes hepatic steatosis by increasing the expression of the fatty acid transporter Cd36 ^34^. Erk1/2 phosphorylation is associated with increased expression of lipogenesis-related proteins ^21,35^. We confirmed that Orm2 is regulated by the Erk1/2–PPARγ pathway *in vivo* and *in vitro*. Notably, we revealed that the loss of Orm2 did not alter Erk1/2 and PPARγ phosphorylation in mice fed with normal diet. However, with HFD, the loss of Orm2 resulted in the activation of Erk1/2 and PPARγ. These results suggest that Orm2 plays a protective role in hepatic steatosis induced by the Erk1/2-PPARγ-Cd36 signaling pathway only under HFD.

Although there is good evidence for the protective role of Orm2, there are some limitations in this study. First, the cause of the high expression of Orm2 in the two lean mice was not explained well. Even though the bioinformatics analysis revealed the Orm2 SNP site variation in population, more genetic studies on humans or mice need to be performed in the future. Second, given that Orm2 is a secreted protein, its multiple effects involve multiple receptors in different tissues. For example, Orm2 binds to Ccr5 on microglia to inhibit LPS-induced neuroinflammation ^16^, Lepr in the hypothalamus to control food intake ^36^, and ITPR2 in hepatocytes ^31^. The deep receptor mechanism of Orm2 and its downstream signaling pathways in the liver, especially in the fatty liver state, deserves further investigation. In addition, even as a glycosylated protein, prokaryotic expressed recombinant ORM2 protein has significant therapeutic effects *in vitro* and *in vivo*; therefore, we speculate that the glycosylation of Orm2 may affect its own secretion but has no effect on its function, which requires further research.

In summary, our study demonstrated that Orm2 plays an important role in protecting liver from pathological damage in NAFLD and NASH, thereby maintaining liver cellular and metabolic homeostasis under dietary challenge or other stress-induced injury. The treatment with recombinant ORM2 protein may also provide novel insights into potential strategies for preventing NAFLD and NASH.

## Materials and Methods

### Animals and animal welfare

Orm2 heterozygote C57BL/6 mice were purchase from Cyagen Biosciences (Guangzhou, China). Male mice were used for all *in vivo* experiments. All animal protocols were approved by the Institutional Animal Care and Use Committee of Northwest A&F University. A constant ambient temperature of 22◻, on a 12 h/12 h light/dark with adlibitum access to standard chow food (Jiangsu Xietong Medicine Bioengineering Co., Ltd., SWS9102, Jiangsu, China) and water.

### Mouse treatment

Male wild-type and Orm2 KO mice were fed a high-fat diet (HFD) (Jiangsu Xietong Medicine Bioengineering Co., Ltd., XTHF60, Jiangsu, China) for 16 weeks to induce NAFLD and fed a methionine choline deficiency (MCD) diet (Jiangsu Xietong Medicine Bioengineering Co., Ltd., XTMCD, Jiangsu, China) for four weeks to induce NASH. For ORM2 protein therapeutic experiments, in the NAFLD model, 8-week-old male mice were fed a HFD diet for six weeks to induce the development of NAFLD, followed by the intraperitoneal (I.P.) injection of Orm2 protein (5.0 mg/kg body weight) or phosphate-buffered saline (PBS) every three days for 10 weeks while continuously on HFD. In the NASH model, 8-week-old male mice were fed with MCD for two weeks to induce the development of NASH, followed by the I.P. injection of Orm2 protein (5.0 mg/kg body weight) or PBS every three days for two weeks, while continuously on MCD.

### Blood biochemical parameters

Serum was separated by the centrifugation of whole blood (3,000 rpm, 8 min). Biochemical parameters, alanine aminotransferase (ALT), aspartate aminotransferase (AST), total cholesterol (TC), glucose, triglycerides (TG), HDL cholesterol, and LDL cholesterol, were determined using an automated biochemical analyzer (Hitachi Ltd., Type 7200-202; Tokyo, Japan) and commercial laboratory tests according to the manufacturer’s instructions.

### Cell culture and stimulation

The human normal liver cell line LO2, 293T cell line, and mouse normal liver cell line AML12 were purchased from the American Type Culture Collection. LO2 cells and 293T cells were grown in Roswell Park Memorial Institute-1640 medium (Hyclone, Logan, U.S.), while AML12 were cultured in DMEM/HIGH GLUCOSE medium (Hyclone, Logan, U.S.), both medias were supplemented with 10% fetal bovine serum (Hyclone, Logan, U.S.). Cells were expanded in standard cell culture conditions: humidified atmosphere, 37◻, 5% CO_2_.

To mimic the hepatic steatosis model *in vivo*, LO2 cells and AML12 cells were treated with cell culture medium containing the indicated concentrations of palmitic acid (PA, P9767; Sigma-Aldrich, MO, USA) and oleic acid (OA, O7501; Sigma-Aldrich, MO, USA) for 24 h; fatty acid-free BSA (B2064; Sigma-Aldrich, MO, USA) was used as a control. PA and OA were used at concentrations of 0.2 and 0.4 mM in LO2 cells and AML12 cells. Recombinant mouse ORM2 were used to stimulate cells (0.5 μg/mL) for OA/PA periods. Pioglitazone were used to stimulate cells (10 μM) for OA/PA periods.

Gene knockdown was performed by introducing (si)RNA of the target gene using Lipofectamine 2000 (Invitrogen, CA, U.S.) following the manufacturer’s instructions. The Orm2 and STAT1 (si)RNA used in the article were purchased from Gene Pharma (Shanghai, China): si-Orm2-sense (5’-3’) GCU UCU AUA ACU CCA GU UAT T; si-Orm2-antisense (5’-3’) UAA CUG GAG UUA UAG AAG CTT; si-STAT1-sense (5’-3’) CAC GAG ACC AAU GGU GUG G; si-STAT1-antisense (5’-3’) CCA CAC CAU UGG UCU CGU G.

The full-length cDNA of Orm2 was amplified from human hepatocyte derived mRNA by RT-qPCR using the following primers: 5′-GAG ATA TCA TGG CGC TGT CCT GG GTT C-3′ and 5′-CCC TCG AGC TAG GAT TCC CCC TCC TCC TG-3′. The gene was cloned into a pcDNA3.1+ vector and assigned as pcDNA3.1-Orm2 recombinant plasmid. The recombinant plasmid pcDNA3.1-Orm2 was transiently transfected into LO2 cells for 48 h with X-treme GENETMHP DNA Transfection Reagent (Roche, Basel, Switzerland).

For luciferase assays, the full-length cDNA of STAT1 was amplified from human hepatocyte derived mRNA by RT-qPCR using the following primers: 5′-GGG GTA CCG CAC AAG GTG GCA GGA TGT C-3′ and 5′-CGG GAT CCT CTA CAG AGC CCA CTA TCC-3′, the Orm2 promoter using the following primers: F1-5′-GGG GTA CCA TAG CTG TTG CCA CAC TC-3′; F2-5′-GGG GTA CCG ACC AAC GTG GAG AAA CC-3′; F3-5′-GGG GTA CCA GCA AGG GAG ACG AGA AG-3′; R1-5′-CCA AGC TTC ATA CTG AGA CCA GGA GG-3′.

### Histological analysis

Liver tissues were fixed in 4% paraformaldehyde, embedded in paraffin, and stained with hematoxylin and eosin (H&E). Picro-Sirius Red staining were carried out to visualize liver fibrosis by service-bio (Hubei, China). Frozen liver sections were subjected to Oil Red O (O9755; Sigma-Aldrich, MO, USA) staining to determine lipid droplet accumulation. Histological images of tissue sections were captured with a light microscope (Nikon eclipse Ni, Tokyo, Japan).

### Immunohistochemistry staining

Tissue sections were incubated with Cd36 antibody in 2% bovine serum albumin-phosphate-buffered saline (blocking solution 4◻, overnight), followed by three times washing and visualization using poly-HRP anti-goat IgG detection kit (ZSGB◻Bio, Beijing, China).

### Real-time quantitative RT-qPCR

RNA extraction, cDNA synthesis, and gene expression measurements of tissues and cells were performed as described previously ^37^. The primer sequences for genes are listed in Supplement Table 1. For RT-qPCR, Gapdh was used to quantify the target gene RNA, by comparative CT method (2^^−ΔΔCt^), as described elsewhere ^38,39^.

### Western blotting

Tissues (50 μg) were homogenized and cells were lysed (lysis buffer, Beyotime, Shanghai, China) containing cocktails and 1% PMSF on ice for 30 min, followed by centrifugation (12, 00 g, 4◻, 15 min). Total protein concentration was determined by a BCA protein assay (Thermo Scientific, MA, U.S.). Proteins were size fractionated by sodium dodecyl sulfate-polyacrylamide gel electrophoresis and transferred into a polyvinylidene difluoride membrane (Roche, Basel, Switzerland). Membranes were incubated (overnight, 4◻) with one of the primary antibodies. After washing TBST 3 times, being incubated with a secondary antibody (2 h, room temperature). Protein bands were visualized by ECL Plus (Thermo Scientific, MA, U.S.). Antibodies used for Western blotting are listed in Supplement Table 2.

### Oral glucose tolerance test and insulin tolerance test

For oral glucose tolerance test, mice were fasted for 16 h and then gavaged with glucose (1.0 g/kg body weight). For insulin tolerance test, mice were fasted for 3 h and injected intraperitoneally with insulin (1.0 U/kg). Blood glucose was measured at 0, 15, 30, 60, 90 and 120 min after gavage or injection by a glucometer.

### Expression and purification of recombinant mouse ORM2

The full-length cDNA fragment of Orm2 was cloned into bacterial expression vector pET-28a, which was then transformed into E. coli BL21 (DE3) strain. The transformed E. coli cells were incubated for 5 h with 1 mM isopropyl-β-D-1-thiogalactopyranoside (IPTG) at 30L to induce mouse ORM2 protein expression. Cells were harvested and lysed using a sonicator (Sonics uibra cell, CT, U.S.). ORM2 in the lysate supernatant was purified with a Ni NTA Beads 6FF gravity column (Smart-Lifesciences, Jiangsu, China). Recombinant mouse ORM2 protein was expressed and purified according to published protocols ^24,40^.

### Luciferase assays

Orm2 promoter activity was determined by luciferase assay using a dual luciferase reporter system (Promega, WI, USA) as described ^41^. 293T cells were transfected using with X-treme GENETMHP DNA Transfection Reagent with the pGL3 basic-Orm2-luciferase plasmid containing the STAT1 promoter and the pRL-SV40 plasmid encoding firefly (Renilla)-luciferase and incubated for 24 h. Luciferase activity was integrated over a 10 s period andmeasured using a luminometer (Victor X Light; MA, U.S.). Results were normalized to the activity of Renilla luciferase. All data are depicted as the mean ± standard deviation (S.D.) of four independent experiments.

### RNA-seq analysis

Total RNA was extracted using Trizol reagent (Thermo Scientific, MA, U.S.). following the manufacturer’s procedure. The total RNA quantity and purity were analysis of Bioanalyzer 2100 and RNA 6000 Nano LabChip Kit (5067-1511, Agilent, CA, U.S.), high-quality RNA samples with RIN number > 7.0 were used to construct sequencing library. Then, we performed the 2×150 bp paired-end sequencing (PE150) on an Illumina Novaseq™ 6000 (LC-Bio Technology CO., Zhejiang, China) following the vendor’s recommended protocol. Differential expressed genes (DEGs) were identified by DESeq2 with two standards: (1) fold change larger than 1.5 and (2) adjusted P values less than 0.05. The data have been submitted to Sequence ReadArchive (SRA) database with PRJNA number: PRJNA879305 and PRJNA883060.

### Statistical analysis

The results are expressed as mean ± S.E.M, and statistical comparisons were performed using Student’s *t*-test between two groups. Statistically significant differences compared to controls are indicated as follows: \**P* <0.05, ** *P* <0.01, *** *P* <0.001 and **** *P* <0.0001; #*P* <0.05, ## *P* <0.01, and ### *P* <0.001, #### *P* <0.0001.

## Abbreviations

Acsl1: acyl-CoA synthetase long chain family member 1
ALT: alanine aminotransferase
Ap2: integrase-type DNA-binding superfamily protein
AST: aspartate aminotransferase
Atgl: Adipose triglyceride lipase
BAT: brown adipose tissue
BCA: bicinchoninic acid
Cpt1ɑ: carnitine palmitoyltransferase 1A
Cxcl10: C-X-C motif chemokine ligand 10
Dgat: diacylglycerol O-acyltransferase
ECL: enhanced chemiluminescence
Erk1/2: mitogen-activated protein kinase
eWAT: epididymal white adipose tissue
Fasn: Fatty Acid Synthase
FFA: free fatty acid
FGF: fibroblast growth factor
HDL: high-density lipoprotein
H&E: hematoxylin and eosin
HFD: high fat diet
IL1β: interleukin 1 beta
IP-ITT: intraperitoneal insulin-tolerance tests
IPTG: isopropyl-β-D-1-thiogalactopyranoside
ITPR2: inositol 1,4,5-trisphosphate receptor type 2
KO: Orm2 knockout mice
LDL: low-density lipoprotein
MCD: methionine and choline-deficient L-amino acid diet
Mcp1: Monocyte chemoattractant protein-1
NAFLD: non-alcoholic fatty liver disease
NASH: non-alcoholic steatohepatitis
NCBI: national center for biotechnology information
OA: oleic acid
O-GTT: oral glucose tolerance
Orm2: Orosomucoid 2
PA: palmitic acid
PMSF: phenylmethanesulfonylfluoride or phenylmethylsulfonyl fluorid
PPAR: peroxisome proliferator activated receptor
RT-qPCR: real time quantitative PCR
SHBG: sex hormone-binding globulin
SMA: smooth muscle
STAT1: Signal Transducer And Activator Of Transcription
TBST: Tris buffered saline with Tween-20
TC: total cholesterol
TG: triglyceride
T2DM: type 2 diabetes mellitus
WT: wild type mice

## Acknowledgments

This study was supported by: National Key Research and Development Program No. 2022ZD0208100, Science Fund for Distinguished Young Scholars of Shaanxi Province No. 2022JC-11.

## Author contributions

Q.S. conceived the idea, supervised the study, analyzed data, and acquired funding. L.L., H.S., J.C. and C.D. performed the experiments. L.L. drafted the manuscript with the help of Q.S., X.Y., H.H. The manuscript was read and approved by all authors.

## Declaration of interests

The authors have declared that no competing interest exists.

## Notes

### Competing Interest Statement

The authors have declared no competing interest.

