## Supplemental Table 1, Table 2; Supplemental Figure 1-5 for "Treatment of non-alcoholic steatohepatitis with recombinant Orosomucoid 2, an acute phase protein that attenuates the Erk1/2-PPARγ-Cd36 signaling pathway"

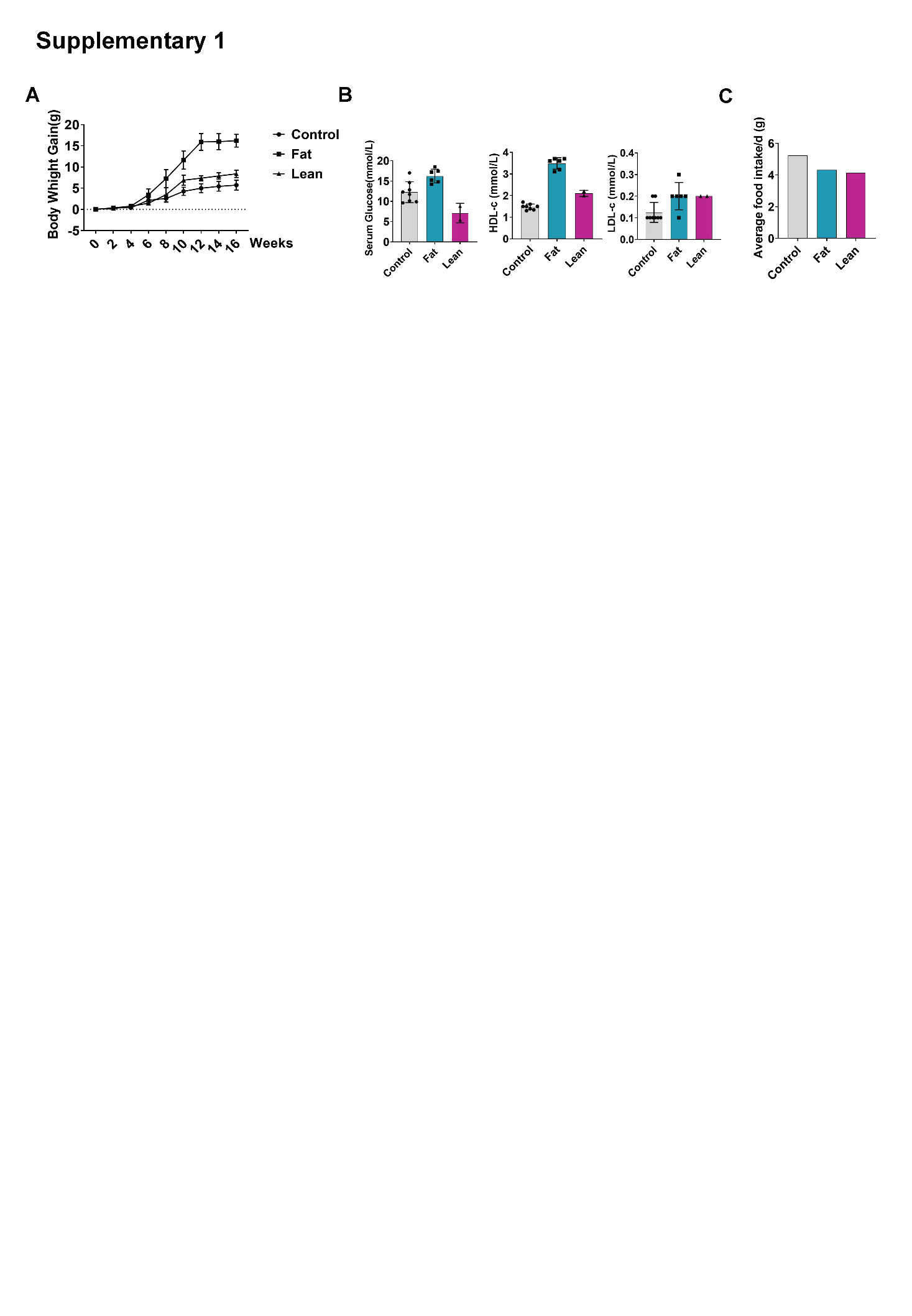
**Supplementary Figure 1** (A) Changes in body weight over sixteen weeks during regular chow (Control), high-fat diet (HFD), and mice fed a high-fat diet but with lower body weight (Lean). (B) Serum glucose, HDL cholesterol, and LDL cholesterol levels. (C) Average daily food intake per mouse over four weeks.

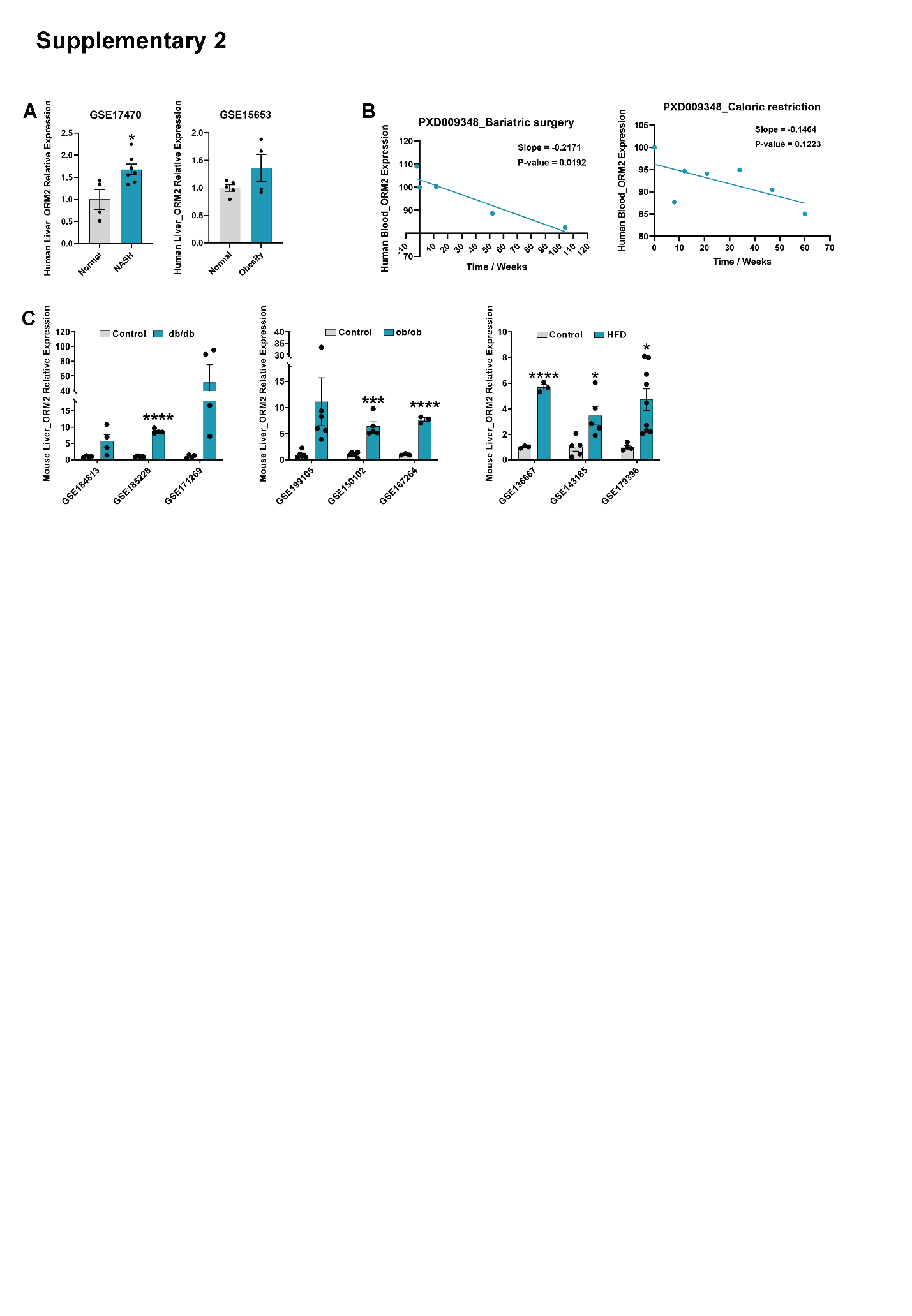
**Supplementary Figure 2** (A) Analysis of ORM2 expression in the human obesity-related liver RNA-seq dataset from the GEO database (<https://www.ncbi.nlm.nih.gov/gds/?term=>). (B) The serum ORM2 content of the PXD009348 dataset in the ProteomeXchange database was analyzed, divided into two parts: bariatric surgery and calorie control, and the trend line was drawn using GraphPad Prism. (C) The expression of ORM2 was analyzed in the RNA-seq datasets of the livers of different obese mouse models in the GEO database. All data are shown as mean±S.E.M. and were compared by Student’s t-test, where **P* < 0.05, ****P* < 0.001, and *****P* < 0.0001.

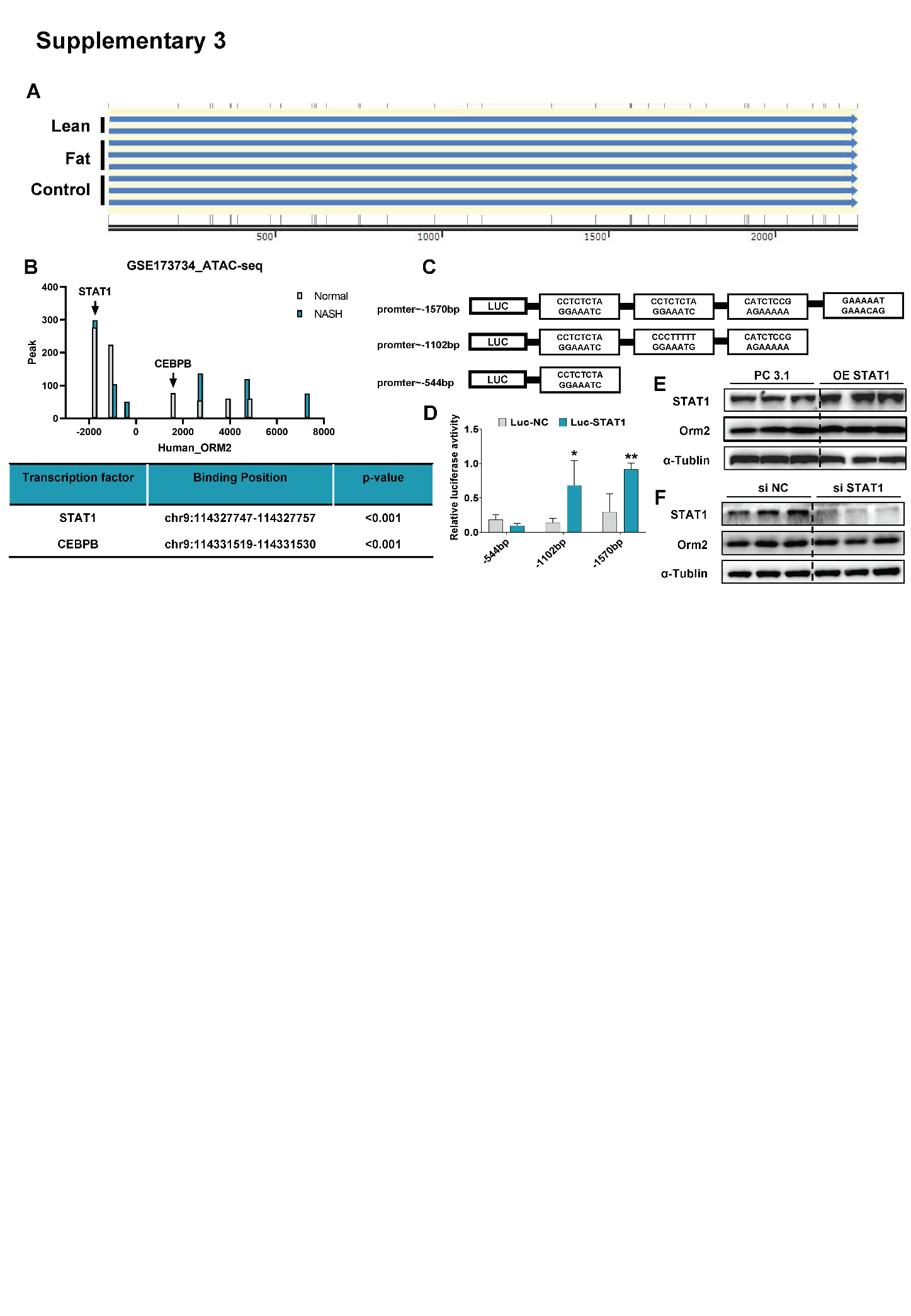
**Supplementary Figure 3** (A) Sequencing and comparison of liver Orm2 promoter regions of Control, Fat, and Thin mice. (B) Statistical analysis of ORM2-related chromatin open regions in the ATAC-seq dataset of GSF173734 in the GEO database, and potential bound transcription factors based on the open sequences using the UCSC Genome Browser website (<http://genome.ucsc.edu/>). predict. (C) Design of a luciferase reporter plasmid to predict the STAT1 binding site in the ORM2 promoter region using the JASPAR database (<http://jaspar.genereg.net/>). (D) Dual luciferase reporter gene, using firefly luciferase for reporter and sea cucumber luciferase for calibration. The reporter plasmid, internal reference plasmid and overexpression plasmid were transferred into HEK-293T cells, and the fluorescence intensity was detected 48 h later. (E) The overexpression plasmid was transferred into LO2 cells. Orm2 and STAT1 were detected by Western Blot after 48 h. (F) The (si)RNA was transferred into LO2 cells. Orm2 and STAT1 were detected by Western Blot after 48 h. All data are shown as mean±S.E.M. and were compared by Student’s t-test, where **P* < 0.05, ***P* < 0.01.

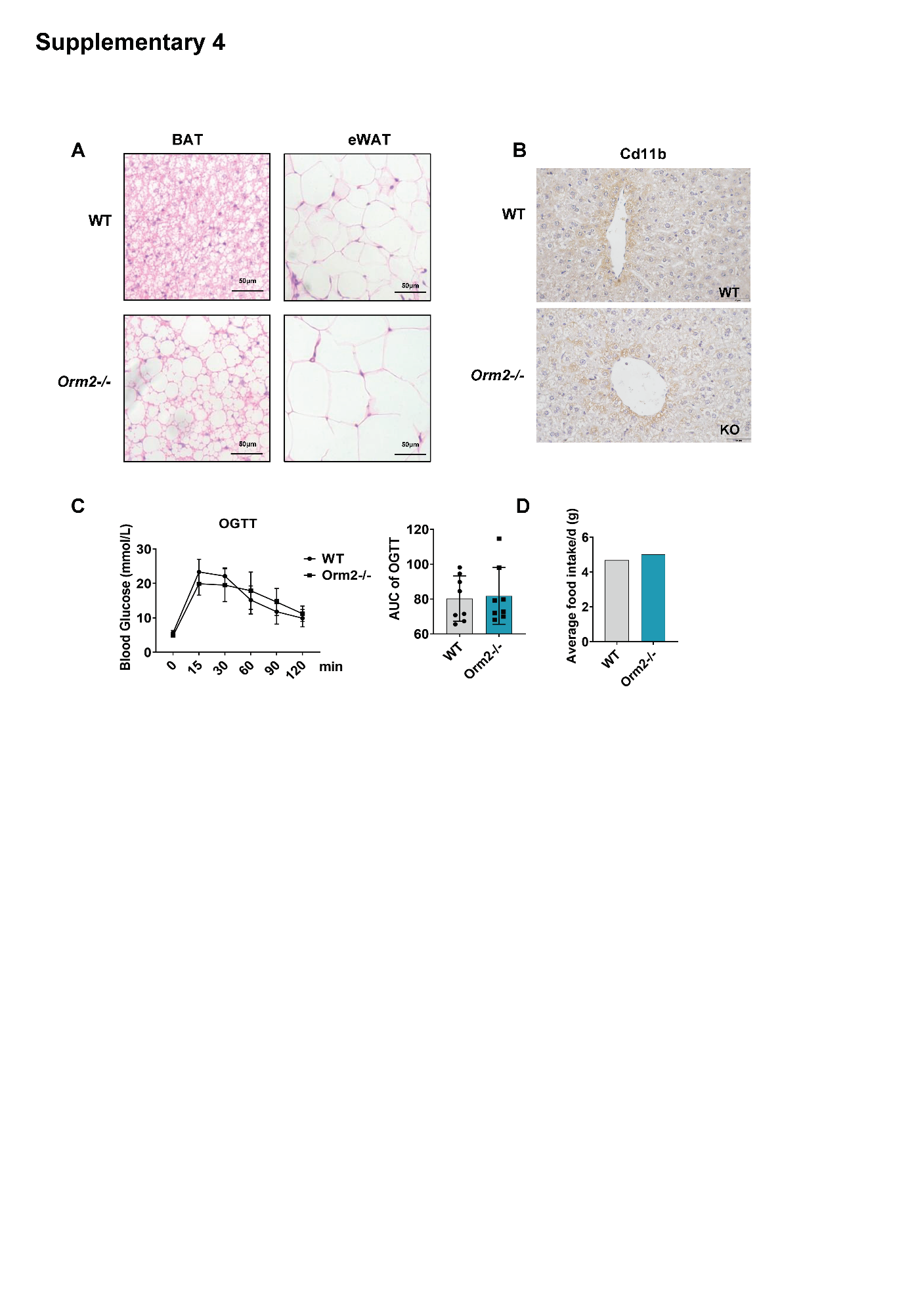

**Supplementary Figure 4** (A) Representative pictures of eWAT and BAT images of H&E staining. Scale bar 50 μm. (B) Representative immunohistochemical image of Cd11b between WT and Orm2-/- mice. (C) Oral glucose tolerance test (OGTT) performed at 17 weeks of age; glucose measurements over time after glucose gavage (left), and area under curve (AUC) shown (right). n = 8 per group. (D) Average daily food intake

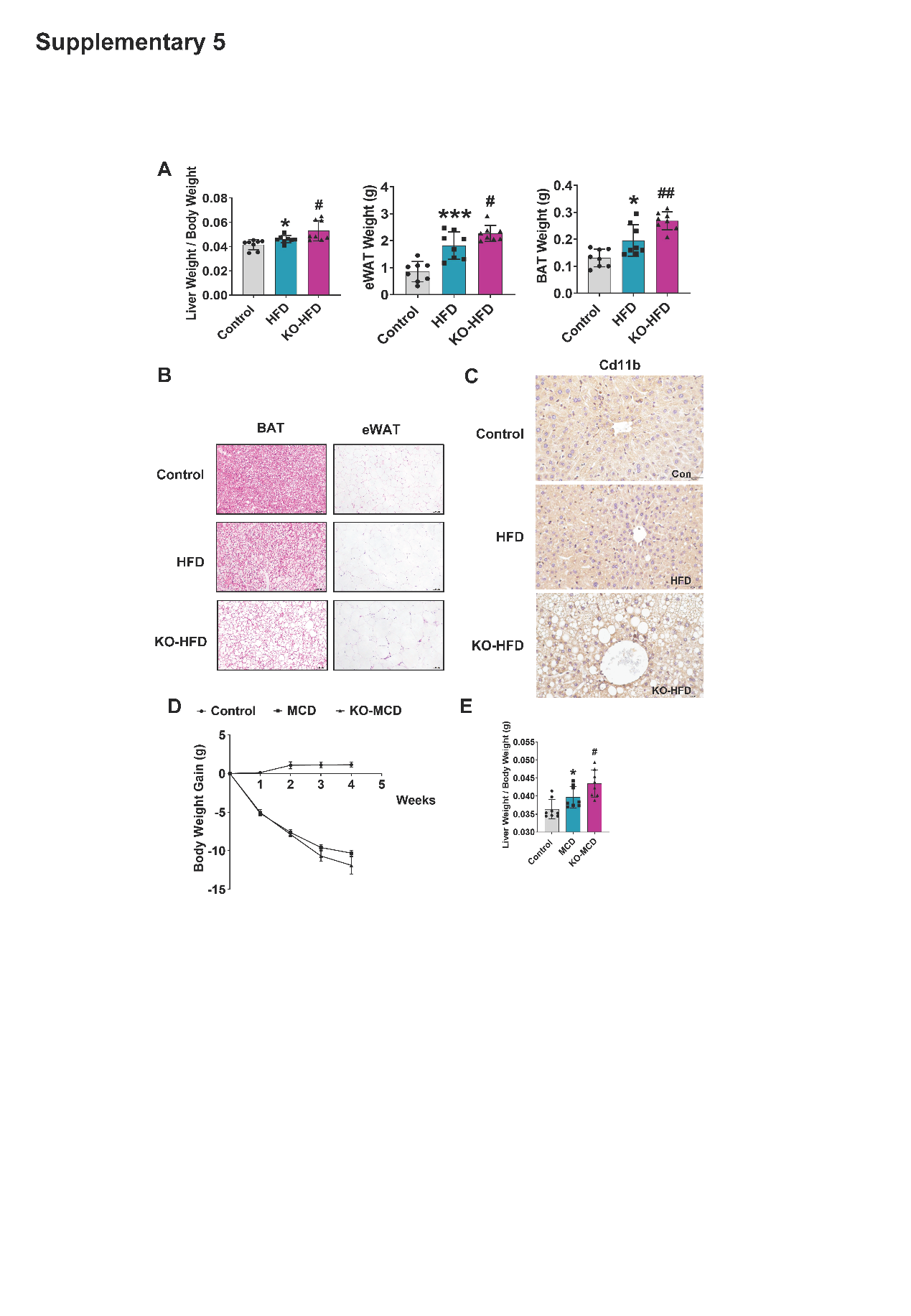
**Supplementary Figure 5** (A) Tissue weights of Liver Weight/Body Weight, BAT and eWAT in mice at 24 weeks of age (n = 8). (B) Representative pictures of eWAT and BAT images of H&E staining. Scale bar 100 μm. (C) Representative immunohistochemical image of Cd11b among Control, HFD, and KO-HFD mice. (D) Body weight change in Control, MCD, and KO-MCD. n =8 per group. (E) Liver Weight/Body Weight in mice at 12 weeks of age among Control, MCD, and KO-MCD, n = 8 per group. All data are shown as mean±S.E.M. and were compared by Student’s t-test. * indicates a significant difference between the Control group and the HFD group; **P* < 0.05, ****P* < 0.001. # indicates a significant difference between the KO-HFD group and the HFD group; #*P* < 0.05, ##*P* < 0.01.

**Supplementary Table 1. Primer sequences for RT-qPCR**

| **Gene Name** | **Sequence (5’-3’)** |
| --- | --- |
| Human-CD36 | Forward: 5’-AAGCCAGGTATTGCAGTTCTTT-3’ |
|  | Reverse: 5’-GCATTTGCTGATGTCTAGCACA-3’ |
| Human-*FATP* | Forward: 5’-GGGGCAGTGTCTCATCTATGG-3’ |
|  | Reverse: 5’-CCGATGTACTGAACCACCGT-3’ |
| Human-*AP2* | Forward: 5’-ACTGGGCCAGGAATTTGACG-3’ |
|  | Reverse: 5’-CTCGTGGAAGTGACGCCTT-3’ |
| Human-*ACC1* | Forward: 5’-ATGTCTGGCTTGCACCTAGTA-3’ |
|  | Reverse: 5’-CCCCAAAGCGAGTAACAAATTCT-3’ |
| Human-*ACC2* | Forward: 5’-AGAAGACAAGAAGCAGGCAAAC-3’ |
|  | Reverse: 5’-GTAGACTCACGAGATGAGCCA-3’ |
| Human-*FASN* | Forward: 5’-GACTGGTACAACGAGCGGAT-3’ |
|  | Reverse: 5’-AGAAGACAAGAAGCAGGCAAAC-3’ |
| Human-*ACSL1* | Forward: 5’-CTTATGGGCTTCGGAGCTTTT-3’ |
|  | Reverse: 5’-CAAGTAGTGCGGATCTTCGTG-3’ |
| Human-*DGAT1* | Forward: 5’-TATTGCGGCCAATGTCTTTGC-3’ |
|  | Reverse: 5’-CACTGGAGTGATAGACTCAACCA-3’ |
| Human-*DGAT2* | Forward: 5’-GAATGGGAGTGGCAATGCTAT-3’ |
|  | Reverse: 5’-CCTCGAAGATCACCTGCTTGT-3’ |
| Human-SCD1 | Forward: 5’-CTTGCGATATGCTGTGGTGC-3’ |
|  | Reverse: 5’-CCGGGGGCTAATGTTCTTGT-3’ |
| Human-*ATGL* | Forward: 5’-GAGAGGGGAGGTTTCCACAC-3’ |
|  | Reverse: 5’-AAGCAGGCGGTCACATACAC-3’ |
| Human-*CPT1α* | Forward: 5’-ATCAATCGGACTCTGGAAACGG-3’ |
|  | Reverse: 5’-TCAGGGAGTAGCGCATGGT-3’ |
| Human-*HSL* | Forward: 5’-CAACTGCCAGCTGCCTTAAA-3’ |
|  | Reverse: 5’-GATTCTGGCTGGGCTATGGG-3’ |
| Human-*GAPDH* | Forward: 5’-GGAGCGAGATCCCTCCAAAAT-3’ |
|  | Reverse: 5’-GGCTGTTGTCATACTTCTCATGG-3’ |
| Mouse-*PPARγ* | Forward: 5’-GGAAGACCACTCGCATTCCTT-3’ |
|  | Reverse: 5’-GTAATCAGCAACCATTGGGTCA-3’ |
| Mouse-*Acsl1* | Forward: 5’-ACCAGCCCTATGAGTGGATTT-3’ |
|  | Reverse: 5’-CAAGGCTTGAACCCCTTCTG-3’ |
| Mouse-*Dgat1* | Forward: 5’-GCCTTACTGGTTGAGTCTATCAC-3’ |
|  | Reverse: 5’-GCACCACAGGTTGACATCC-3’ |
| Mouse-*Dgat2* | Forward: 5’-GCGCTACTTCCGAGACTACTT-3’ |
|  | Reverse: 5’-GGGCCTTATGCCAGGAAACT-3’ |
| Mouse-*Cd36* | Forward: 5’-ATGGGCTGTGATCGGAACTG-3’ |
|  | Reverse: 5’-TTTGCCACGTCATCTGGGTTT-3’ |
| Mouse-*Ap2* | Forward: 5’-AAGGTGAAGAGCATCATAACCCT-3’ |
|  | Reverse: 5’-TCACGCCTTTCATAACACATTCC-3’ |
| Mouse-*Fasn* | Forward: 5’-CTGCCTTCGGTTCAGTCTCTT-3’ |
|  | Reverse: 5’-AGGCCACTTGTGGGGAATAC-3’ |
| Mouse-*PPARα* | Forward: 5’-AACATCGAGTGTCGAATATGTGG-3’ |
|  | Reverse: 5’-CCGAATAGTTCGCCGAAAGAA-3’ |
| Mouse-*Cpt1α* | Forward: 5’-GACTCCGCTCGCTCATTCC-3’ |
|  | Reverse: 5’-ACCAGTGATGATGCCATTCTTG-3’ |
| Mouse-*Mcp1* | Forward: 5’-TAAAAACCTGGATCGGAACCAAA-3’ |
|  | Reverse: 5’-GCATTAGCTTCAGATTTACGGGT-3’ |
| Mouse-*Il1β* | Forward: 5’-GAAATGCCACCTTTTGACAGTG-3’ |
|  | Reverse: 5’-TGGATGCTCTCATCAGGACAG-3’ |
| Mouse-*Cxcl10* | Forward: 5’-CCAAGTGCTGCCGTCATTTTC-3’ |
|  | Reverse: 5’-GGCTCGCAGGGATGATTTCAA-3’ |
| Mouse-*Gapdh* | Forward: 5’-GTTGTCTCCTGCGACTTCA-3’ |
|  | Reverse: 5’-TGGTCCAGGGTTTCTTACTC-3’ |

**Supplementary Table 2. Antibody information**

| **Antibody** | **Source** | **Identifier** | **Application** |
| --- | --- | --- | --- |
| FASN Polyclonal antibody | Proteintech | Cat No.#10624-2-AP | WB (1:1000) |
| ATGL Polyclonal antibody | Proteintech | Cat No : 55190-1-AP | WB (1:1000) |
| Orosomucoid 2 Polyclonal antibody | Abcam | Cat# ab231906; | WB (1:1000) |
| Alpha Tubulin Polyclonal antibody | Proteintech | Cat No.#11224-1-AP | WB (1:1000) |
| Collagen Ⅰ polyclonal antibody | Wanleibio | Cat# WL0088 | WB (1:500) |
| alpha-SMA Polyclonal Antibody | Affinity | Cat# AF1032 | WB (1:1000) |
| Phospho-PPAR gamma (Ser112) Antibody | Affinity | Cat# AF3284 | WB (1:1000) |
| PPAR gamma Polyclonal Antibody | Affinity | Cat# AF6284 | WB (1:1000) |
| Phospho Erk1/2 (Thr202/Tyr204) Rabbit mAb | Cell Signaling Technology | Cat#4370 | WB (1:1000) |
| Erk1/2 Rabbit mAb | Cell Signaling Technology | Cat#4695 | WB (1:1000) |
| Mouse monoclonal anti-CD36 | Santa Cruz Biotechnology | Cat# sc-7309 | WB (1:200) |
| CD11b/ITGAM Rabbit pAb | ABclonal | Cat#A1581 | IHC (1:100) |
| Peroxidase-Conjugated Goat Anti-Mouse IgG (H+L) | Diyibio | Cat#DY60203 | WB (1:2000) |
| Peroxidase AffiniPure Goat Anti-Rabbit IgG (H+L) | Diyibio | Cat#DY60202 | WB (1:2000) |
